# Modeling Antigen-Specific T Cell Dynamics Following Hepatitis B Vaccination indicates differences between conventional and regulatory T cell dynamics

**DOI:** 10.1101/2023.05.27.542434

**Authors:** Hajar Besbassi, George Elias, Pieter Meysman, Hilde Jansens, Kris Laukens, Pierre Van Damme, Niel Hens, Philippe Beutels, Benson Ogunjimi

## Abstract

Our study aims to investigate the dynamics of conventional memory T cells (Tconv) and regulatory T cells (Treg) following activation, and to explore potential differences between these two cell types. To achieve this, we developed advanced statistical mixed models based on mathematical models of ordinary differential equations (ODE), which allowed us to transform post-vaccination immunological processes into mathematical formulas. These models were applied on in-house data from a de novo Hepatitis B vaccination trial. By accounting for inter- and intra-individual variability, our models provided good fits for both antigen-specific Tconv and Treg cells, overcoming the challenge of studying these complex processes. Our modeling approach provided a deeper understanding of the immunological processes underlying T cell development after vaccination. Specifically, our analysis revealed several important findings regarding the dynamics of Tconv and Treg cells, as well as their relationship to seropositivity for HSV-1 and EBV, and the dynamics of antibody response to vaccination. Firstly, our modeling indicated that Tconv dynamics suggest the existence of two T cell types, in contrast to Treg dynamics where only one T cell type is predicted. Secondly, we found that individuals who converted to a positive antibody response to the vaccine earlier had lower decay rates for both Tregs and Tconv cells, which may have important implications for the development of more effective vaccination strategies. Additionally, our modeling showed that HSV-1 seropositivity negatively influenced Tconv cell expansion after the second vaccination, while EBV seropositivity was associated with higher Treg expansion rates after vaccination. Overall, this study provides a critical foundation for understanding the dynamic processes underlying T cell development after vaccination.

## 1 Introduction

The immune system plays a crucial role in protecting the body against pathogens and preventing the development of diseases. It is a complex system consisting of various types of cells and molecules that work together to identify and eliminate foreign invaders while maintaining self-tolerance. Among the key players of the immune system are T cells, which play a central role in orchestrating immune responses against pathogens and cancer cells^1, 2^. Memory T cells, both conventional and regulatory, are subtypes of T cells that provide long-lasting protection against previously encountered antigens and modulate immune responses, respectively^3, 4^. Conventional memory T cells are responsible for the rapid and robust response to pathogens upon re-exposure, while regulatory T cells suppress immune responses and prevent excessive inflammation and tissue damage^3, 4^. The development and maintenance of memory T cells are complex processes that involve various factors such as cytokines, co-stimulatory molecules, and transcription factors^5, 6^.

Vaccination is a highly effective strategy for generating memory T cells and protecting individuals from infectious diseases^7^. For example, the hepatitis B virus (HBV) vaccine induces the production of memory T cells and antibodies against the virus, providing long-term protection against HBV infection^8^. Anti-hepatitis B surface antibodies (Anti-HBs) are proteins produced by the immune system in response to the presence of the hepatitis B surface antigen (HBsAg). Anti-HBs concentrations more than 10 mIU/mL provide protection against infection. However, it is worth noting that the percentage of individuals maintaining anti-HBs levels above 10 mIU/mL is significantly influenced by the duration since the initial vaccination. Anti-HBs concentrations below this threshold may not confer the same level of protection against HBV infection. It is important to recognize that the 10 mIU/mL threshold is a commonly used indicator of protection, although it is not an absolute cutoff. Memory specific T cells prompt a powerful anamnestic response upon exposure to the hepatitis B virus, preventing acute illness, long-term viremia, and chronic infection. An anamnestic anti-HBs response following vaccination with an additional dose demonstrates specific memory T cells after hepatitis B vaccination^9^. The majority of individuals who had a favorable response to the initial series of vaccinations will experience a rapid rise in anti-HBs as a result of this booster dose. Time since first vaccination appears to be related to immune memory persistence^10^. Although progress has been made in the development of vaccines against hepatitis B virus (HBV), the long-term clinical outcome of the disease remains poor due to the challenge of achieving immunological memory, which may involve viral clearance and/or non-specific antibody response^11, 12^.

Recently, several mathematical models were developed to model longitudinal immune responses using ordinary differential equations (ODEs), which are popular and powerful tools for modelling very complex dynamical systems in many fields. They are widely used in the study of population dynamics, epidemiology, virology, pharmacokinetics and gene regulatory networks because of their ability to describe key interaction mechanisms between biological components of complex systems, their evolution over time, and provide reasonable stochastic dynamics approximations^13–17^. Mixed-effects modeling offers the benefit of accounting for and quantifying the correlation between several replicates, as well as of achieving a more precise parameter estimate by pooling all the data, in comparison to fixed-effects modelling. There is currently a growing interest in estimating mixed-effects ODE models due to their ability to account for both within- and between-subject variability. With repeated measurements from multiple individuals, mixed-effects ODE models provide a more robust way of estimating model parameters than traditional ODE models. As a result, there is a growing interest in developing and applying mixed-effects ODE models in immunology research. The first to use the mixed effects modeling method with ordinary differential equations (ODE) in a form that more closely approximated immune response dynamics after vaccination were Andraud et al.^16^, who focused on long-term impacts, and Le et al.^18^ on the immediate effects of vaccination. Furthermore, Keersmaekers et al.^19^ used the mixed effect modeling to investigate two vaccine doses. Another recent study by Besbassi et al.^20^ employed mixed-effects ODE models to examine antibody dynamics following re-exposure to infection. The approach was applied to 61 herpes zoster patients to gain insights into varicella-zoster virus specific antibody dynamics during and after clinical herpes zoster. The study provided a deeper understanding of the population’s characteristics and offered unique insights that can aid in improving our understanding of VZV antibody dynamics and in making more accurate projections regarding the potential impact of vaccines^21^.

In this current study, we applied an ODE-based mixed effects modeling approach to data related to the kinetics of hepatitis B virus (HBV)-specific memory T cell responses in individuals who received de novo HBV vaccination^12^. Our aim was to characterize the dynamics of memory T cell responses over time and identify factors that influence the development and maintenance of immunological memory. Overall, our study highlights the complex dynamics of memory conventional and regulatory T cell responses and the importance of considering individual variability in the development and maintenance of immunological memory. Our findings may have important implications for the design of vaccination strategies and the development of new immunotherapies for infectious diseases.

## 2 Material and methods

### 2.1 Data

We used data from a previously published study by Elias et al.^12^, which included a cohort of 34 healthy subjects who received the hepatitis B vaccine. The participants had no history of HBV infection or previous hepatitis B vaccination. The vaccine used was Engerix-B with a dose of 20 *μg* of hepatitis B surface antigen with alum adjuvant, administered by intramuscular injection on days 0, 30, and 365. Peripheral blood samples were collected at four time points: day 0 (pre-vaccination), as well as 3 months, 6 months, and 12 months after the first vaccine dose. The subjects were classified into three groups based on their antibody response to the vaccine: early responders, late responders, and non-converters. Early responders were defined as individuals with an anti-HBs antibody titer ≥ 100 IU/L at month 3, late responders were defined as individuals with an anti-HBs antibody titer ≥ 100 IU/L at month 6 or month 12, and non-converters were defined as individuals with an anti-HBs antibody titer *<* 10 IU/L at month 12.

In addition to age and sex, the study also collected data on the TCR affinity for the HBsAg peptide pool and cytomegalovirus (CMV), ebstein-barr virus (EBV), or herpes simplex virus type 1 (HSV) seropositivity during the study. The TCR affinity, specifically the HBsAg-specific TCR affinity, was determined by calculating the ratio of unique T-cell receptors (TCRs) annotated as HBsAg-specific in the sequenced TCR repertoire to the number of bystander TCRs. This calculation provides insight into the strength of the binding interaction between the T-cell receptor and the peptides derived from the hepatitis B surface antigen. CMV, EBV, and HSV seropositivity were measured to evaluate whether previous exposure to these viruses influenced the immune response to the hepatitis B vaccine. The data collected was used to study the longitudinal dynamics of CD4+ T cells and to evaluate the influence of these factors on the immune response to the hepatitis B vaccine.

### 2.2 T-cell data

Generating adaptive immune responses against microbial invaders is mostly dependent on CD4+ T cells. The identification of antigenic peptides presented on major histocompatibility complexes (MHC) by the TCR, together with antigen-independent co-stimulation, is necessary for naive CD4+ T cells to develop into more specialized subsets after T cell activation. Depending on the antigen, CD4+ T cells can differentiate into a variety of subset populations. In the previously published study^12^ HbsAg stimulated CD4+ T cells were subtyped into different cell subsets; however, we will only focus on two types: one being the memory conventional T cell subset out of total CD4+ T cell (“Tconv”), which is defined as CD154+CD137-cells. The other being regulatory memory T cells out of total CD4+ T cells (“Tregs”), which is defined as CD154-CD137+ cells, that specialise in immunological homeostasis and maintenance of self-tolerance, inflammation control and prevention of autoimmune diseases.

Figures 1 and 3 show individual-specific memory Tconv and Treg profiles by time in days, respectively. The data are presented for early, late, and non-converters. These figures are used to analyze the explore of age, gender, anti-HBs antibody titers, TCR affinity for the HBsAg peptide pool, and CMV, EBV, or HSV seropositivity on longitudinal CD4+ T cell dynamics shown in Figures 2 and 4.

**Figure 1.**
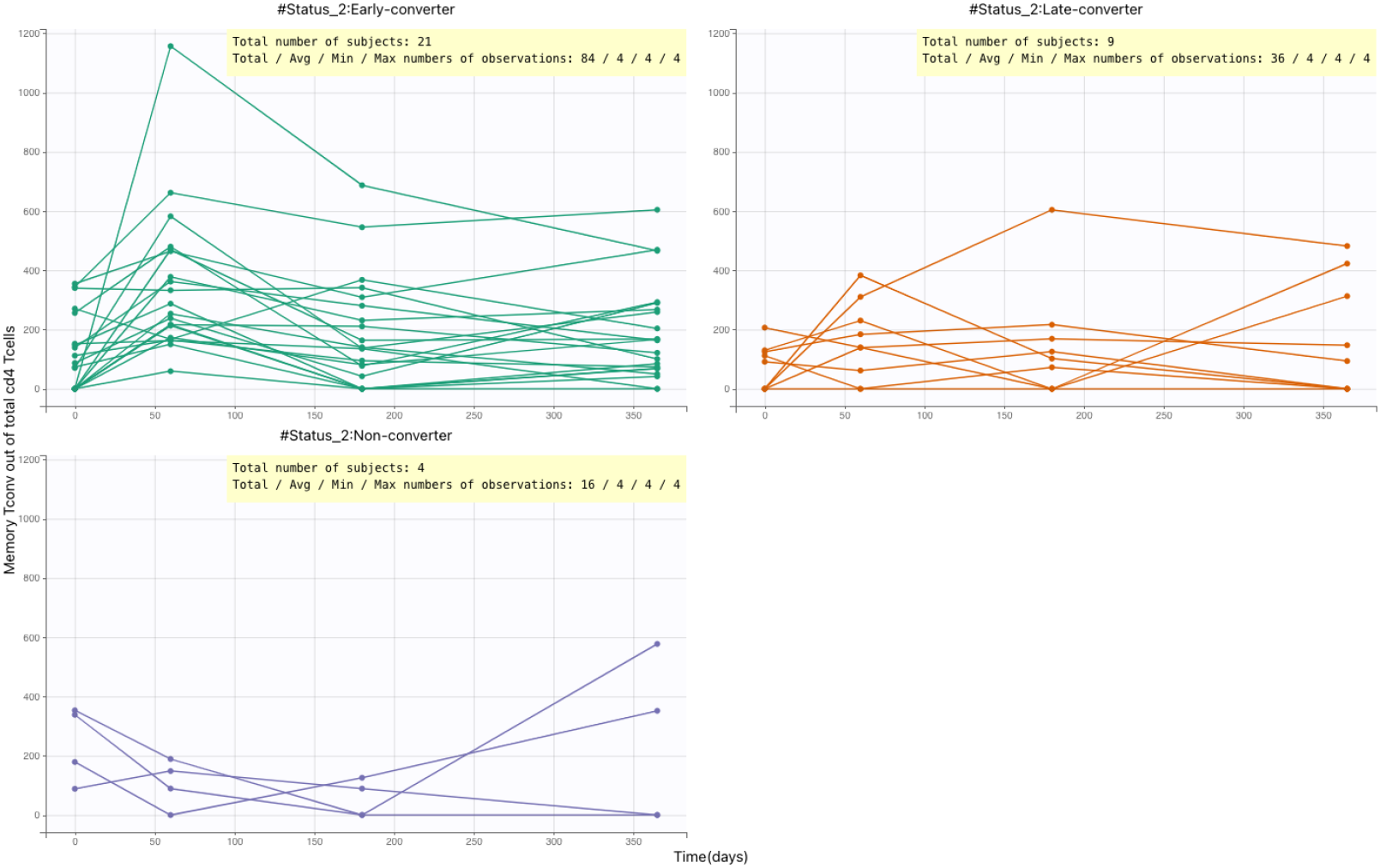
Individual-specific memory Tconv profiles by time (in days) for Early/Late/Non converter status. The plot shows the variation in memory Tconv over time for each individual, and how this varies between early converters, late converters, and non-converters.

**Figure 2.**
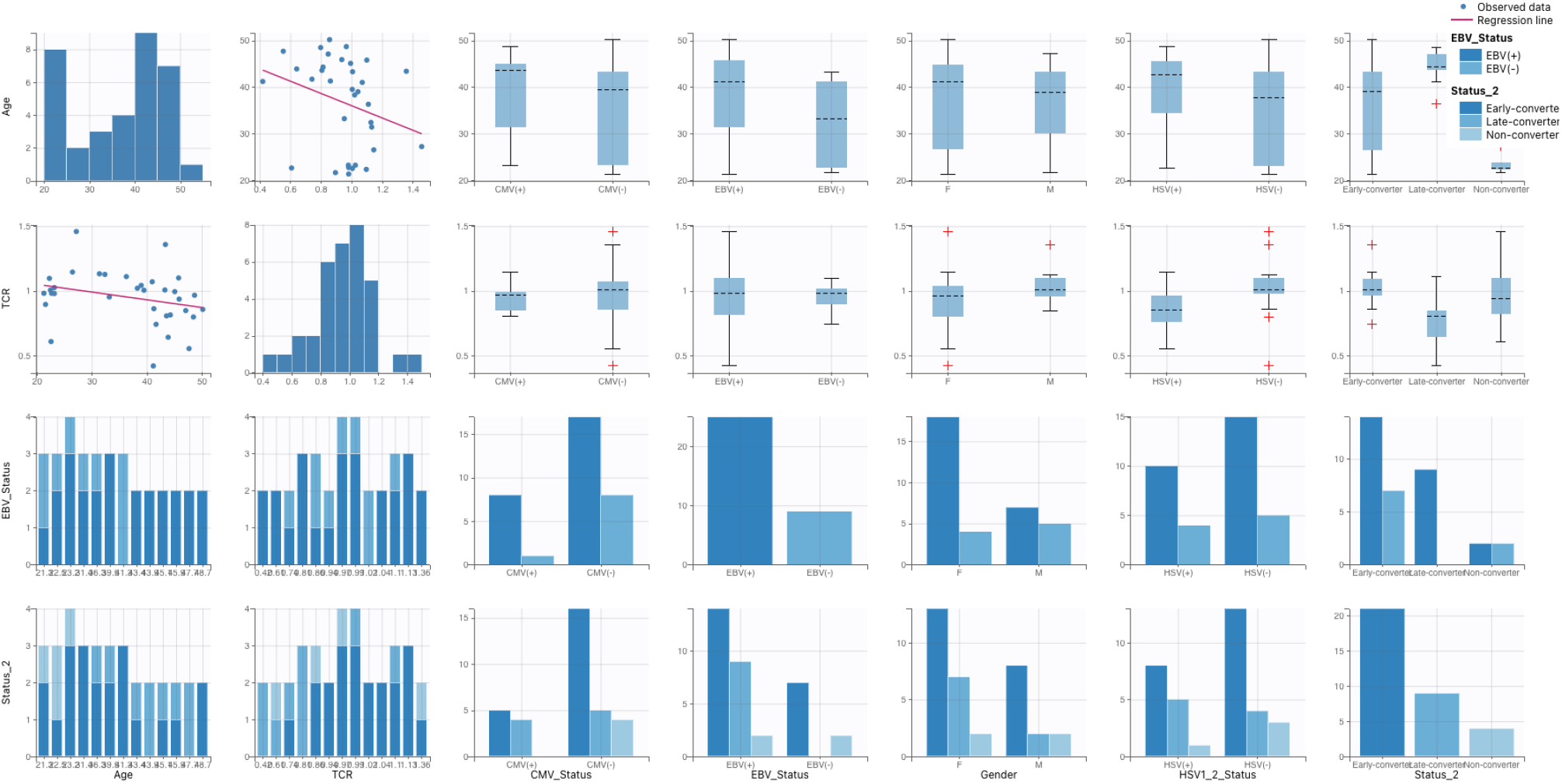
Matrix of Significant Covariates on Tconv: Effect of continuous covariates Age, TCR, and Categorical covariates CMV, EBV, HSV1-2, Converter Status. The figure illustrates the impact of each covariate on the outcome and allows for a comparison of their effect sizes.

**Figure 3.**
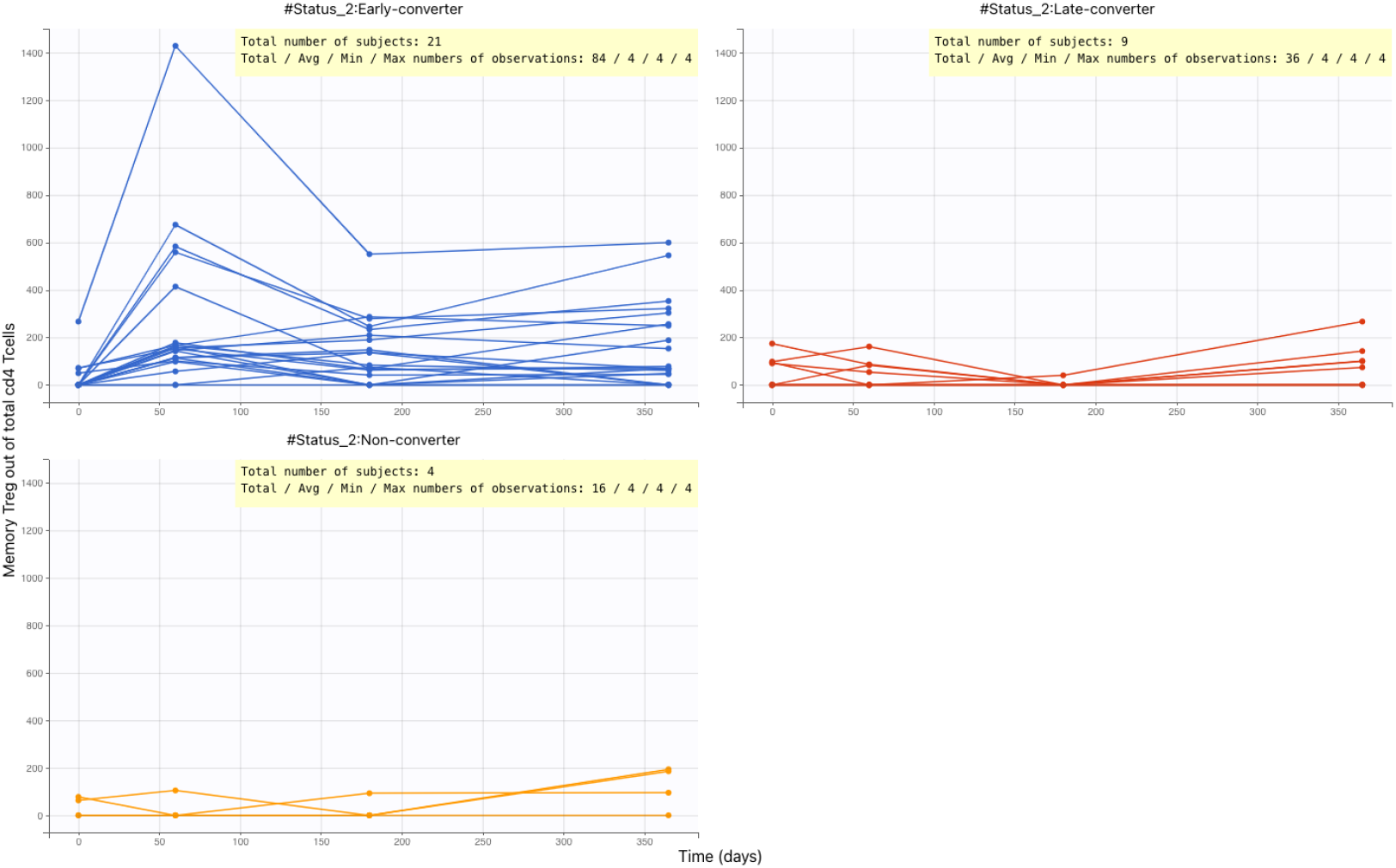
Longitudinal profiles of memory regulatory T-cells over time (in days) for individual participants, stratified by their converter status (Early/Late/Non). The variation in response over time and between individuals is apparent, highlighting the importance of individual-level modeling approaches

**Figure 4.**
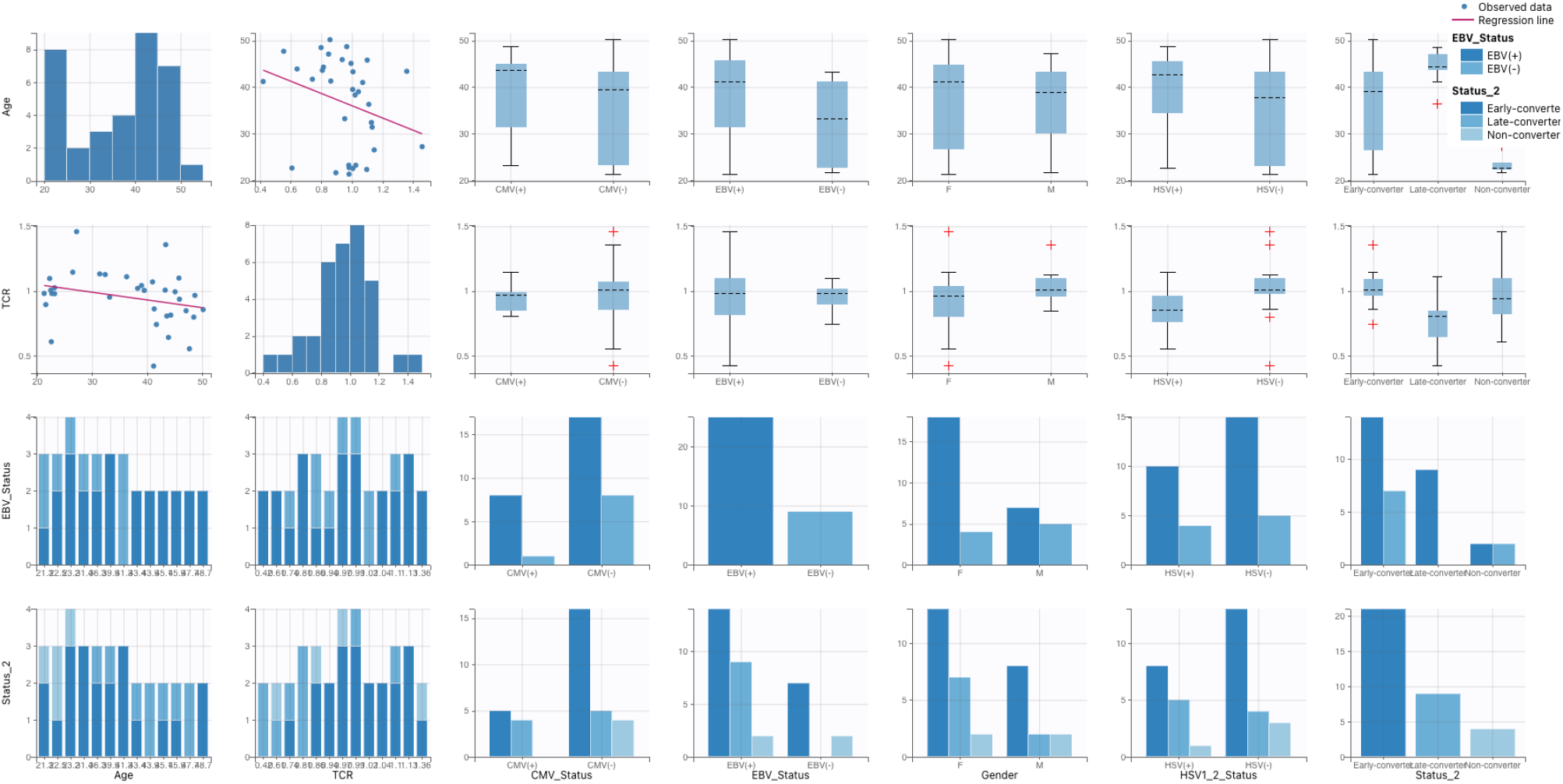
Matrix of Significant Covariates on Treg: Effect of continuous covariates Age, TCR, and Categorical covariates CMV, EBV, HSV1-2, Converter Status. The figure illustrates the impact of each covariate on the outcome and allows for a comparison of their effect sizes.

### 2.3 Mathematical methods

#### 2.3.1 T-cell dynamic Models

In this subsection, we present nonlinear mixed-effect models based on ordinary differential equations to model the dynamics of T-cells. Our goal is to obtain models that best describe the available data while representing the important cell populations in the T-cell response process generated by the body after vaccination. The main T-cell populations considered include conventional memory T cells, regulatory memory T cells. These are further subdivided into short-lived and long-lived memory T conventional cells, and short-lived and long-lived memory T regulatory cells.

We used a systematic approach to fit and compare multiple models to obtain the least number of parameters needed to accurately describe the T-cell dynamics. The general ODE equation used to describe the dynamics of T-cells is:

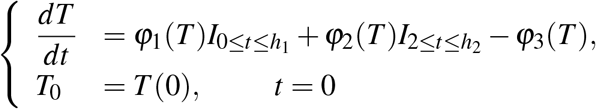

Here, *φ*_1_ represents the expansion of T-cells after the first vaccination at time 0 until a certain time *h*_1_ (with 0 ≤ *h*_1_ ≤ 2). After *h*_1_ has been reached, T-cells will not further be activated until the second vaccination, one month after the first one, which *φ*_2_ defines as the expansion of T-cells during the period [2, *h*_2_], *h*_2_ being the point at which the second T-cell peak is attained. The decay of T-cells will occur throughout the full time period and is described by the function *φ*_3_. In all models, we assume that the decay rate of T-cells remains constant and is proportional to the number of T-cells, which can be written as *φ*_3_ = *μ*_*T*_ *× T*. Moreover, we assume a constant expansion rate for T-cells after each vaccination event, which aligns with the initial phase of the immune response characterized by rapid T-cell activation and subsequent division. In this model, one can hypothesize that 1 dividing T-cell will generate 1 circulating “effector” T-cell and 1 T-cell that will proceed in the expansion process.

Assuming an equal and constant expansion rate of T cells after each vaccination leads to Model 1. Functions *φ*_*i*_(*i* ∈ {1, 2, 3}) can now be written as:

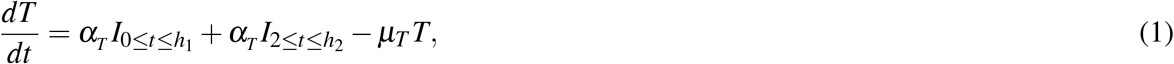

Model 2 does not assume an equal expansion rate after each vaccination. In addition, it is reasonable to assume a different rate after the second vaccination due to a memory response. In this scenario, the functions *φ*_*i*_ are expressed as follows:

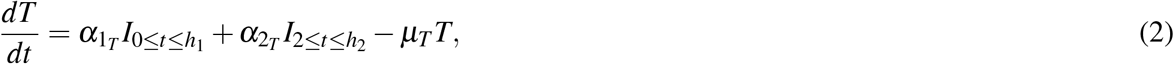

As we proceed, we distinguish the short-lived T cells, denoted as ST(t), from the long-lived T cells, denoted as LT(t). The total T-cell population is then represented as the sum of these two distinct sub-populations. It is important to note that we initially consider only the presence of long-lived T cells and exclude any short-lived T cells at the beginning. This modeling approach enables us to effectively describe the dynamic behavior of T-cell populations during various stages of the immune response. We describe it by the functions *φ*_*i*_, *ψ*_*i*_ where *i* ∈ {1, 2, 3}):

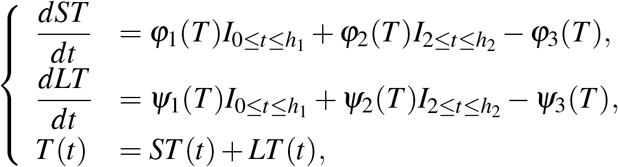

with *LT*_0_ = *LT* (0) the initial number of long living T-cells and *ST*_0_ = 0 the number of short living T-cells at time 0.

First, similarly to the preceding models, we will suppose that the expansion rates of ST are constant and there is no decay of LT (thus a constant number of LT). Model 3 also presumes that the expansion rates of ST after the two vaccinations are identical. The functions *φ*_*i*_ and *ψ*_*i*_ are accordingly given as:

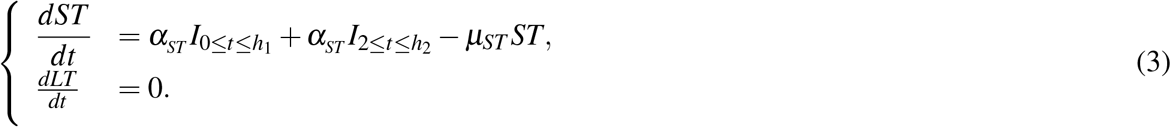

In Model 4, we consider different expansion rates of ST after the two vaccinations, which leads to:

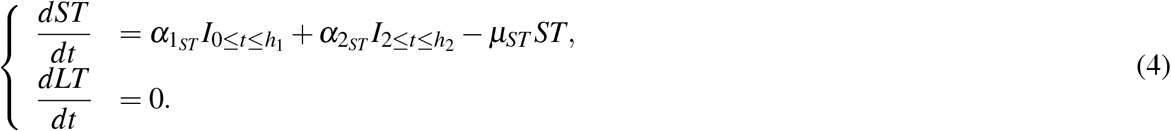

Thereafter, we also introduce a constant *a*_*LT*_ proliferation rate of long-lived T cells after each vaccination. Taking this into account, we obtain Model 5 where we assume equal expansion rates for ST after the two vaccinations, written as :

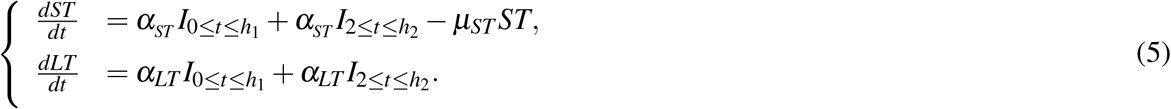

We complete the T cell models with the Model 6, in which in each vaccination different ST activation rates are considered, given as:

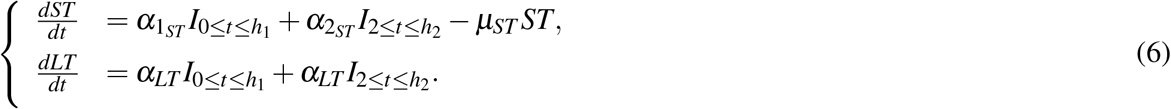

### 2.4 Statistics

#### 2.4.1 Nonlinear mixed models

Nonlinear mixed models can be developed based on the mathematical models discussed in the previous subsection. In this type of model, each individual parameter *P*_*i*_ can be described as *P*_*i*_ = *u*_*i*_*P*_*pop*_, where *P*_*pop*_ is a population parameter and *u*_*i*_ is log-normally distributed with *E*(*u*_*i*_) = 1. To account for categorical variables such as sex, a dummy variable is introduced with an additional parameter *β*_*j*_, which explains how the parameter in group *j* differs from the reference group. This allows for investigating which specific parameter of the structural model, such as expansion rate or decay rate, is responsible for observed differences between different groups. The algorithm used here involves an integrated stochastic approximation of the standard expectation maximization algorithm (SAEM) with simulated annealing, combined with a Markov chain Monte Carlo (MCMC) approach that substitutes the simulation step of the SAEM algorithm. Afterwards, the log-likelihood is computed through importance sampling, where a fixed t-distribution with 5 degrees of freedom is presumed. The data analysis and statistical modeling for this study were conducted using VSC (Vlaams Supercomputer Centrum), Flanders’ highly integrated high-performance research computing environment.

#### 2.4.2 Inference and model selection

The following procedure was used to compare and select the most appropriate biologically credible model to fit the data. In a first step, a list of models was composed, consisting of models 1 to 6 for T-cell data, together with assumptions on the parameters reflecting whether or not individual variation on these parameters is present, i.e. whether or not random effects were included for the different parameters. The model parameters were then estimated with the Monolix software ©Lixoft.

Models with poor SAEM convergence, likely because of abundant model complexity, were discarded. Next, the candidate models were compared using Akaike’s Information Criterion (AIC) and the model with lowest AIC value was selected as first candidate model. Subsequently, a non-parametric bootstrap, using 1000 bootstrap re-samples, was performed on the candidate model. Since a sequential approach based on the candidate models with the lowest AIC values was used, the need to perform bootstraps for all candidate models was avoided, in order to decrease the number of computations. It was found that for a bootstrap, either 70-80% of the samples had proper SAEM convergence. For this reason the criterion for good bootstrap convergence was defined as having at least 70% of bootstrap samples with proper SAEM convergence. In case of poor bootstrap convergence, the candidate model was rejected from the list of candidate models. Please see^14^ for additional details on the algorithms used. The algorithms mainly used default values for Monolix parameters. The convergence of the estimate of population parameters is assessed by the two-step SAEM-MCMC technique.

## 3 Results

For the present study, all parameters were initially set as random and were later selected one by one to be fixed. After testing each parameter individually, all parameters were ultimately fixed. The selection of the fixed effects was based on a thorough analysis of their statistical and theoretical significance. Only the significant fixed effects were included in the table 1 and table 3, which was generated through a series of preliminary analyses that involved running multiple models with different combinations of fixed and random parameters. The rigorous selection process helped to ensure that the fixed effects included in the final tables were meaningful and contributed to the overall scientific rigor of the study.

**Table 1.** ODE Model Formulations Considered for Memory Conventional T-Cell Data and Model Selection Procedure. The table shows the fixed population parameter, convergence-AIC, candidate models, and bootstrap convergence for the six different Tconv models considered in the analysis. Model 4c and 6a were selected as the best models for the data, with 91% and 73% bootstrap convergence, respectively. The table provides important information about the model selection procedure for the Tconv models.

| Tconv model | Fixed population parameter | Convergence-AIC | Candidate model | Bootstrap convergence |
| --- | --- | --- | --- | --- |
| 1a | — | 1727,23 | No |  |
| 2a | — | 1705,11 | No |  |
| 2b | $h_2$ | 1701,86 | No | |
| 3a | — | 1756,3 | No |  |
| 3b | $\mu_{ST}$ | No | No | |
| 4a | — | 1651,33 | Yes | 91% |
| 4b | $\alpha_{1ST}$ | No | No | |
| 4c | $\alpha_{2ST}$ | 1649,52 | Yes | No |
| 4d | $\mu_{ST}$ | No | No | |
| 4e | $h_1$ | No | No | |
| 4f | $h_2$ | No | No | |
| 5a | — | 1760,13 | No |  |
| 5b | $h_2$ | 1758,05 | No | |
| 6a | — | 1660,01 | Yes | 73% |
| 6e | $h_1$ | 1656,23 | Yes | No |

**Table 2.** Parameter estimates and corresponding 95% confidence intervals (CI) of final model 4a.

| Parameter | Estimate | 95% CI | P-value |
| --- | --- | --- | --- |
| $T_{conv}(0)$ | 10976.43 | (6933096 ; 897741604180 ) | |
| $\alpha_{1_{ST}}$ | 10.65 | ( 7.62; 31.689) | |
| $\alpha_{2_{ST}}$ | 3.13 | ( 0.615 ; 7.053) | |
| $\beta_{\alpha_{2_{ST}}}(HSV1)$ | -1.65 | (-2.918; -0.292) | 4.63e-16 |
| $u_{ST}$ | 0.017 | (0.006; 0.045) | |
| $\beta_{u_{ST}}(Late - converter)$ | 1.28 | (0.18; 2.336) | 3.04e-6 |
| $\beta_{u_{ST}}(Non - converter)$ | 1.92 | (0.719; 3.271) | 4.08e-7 |

**Table 3.**
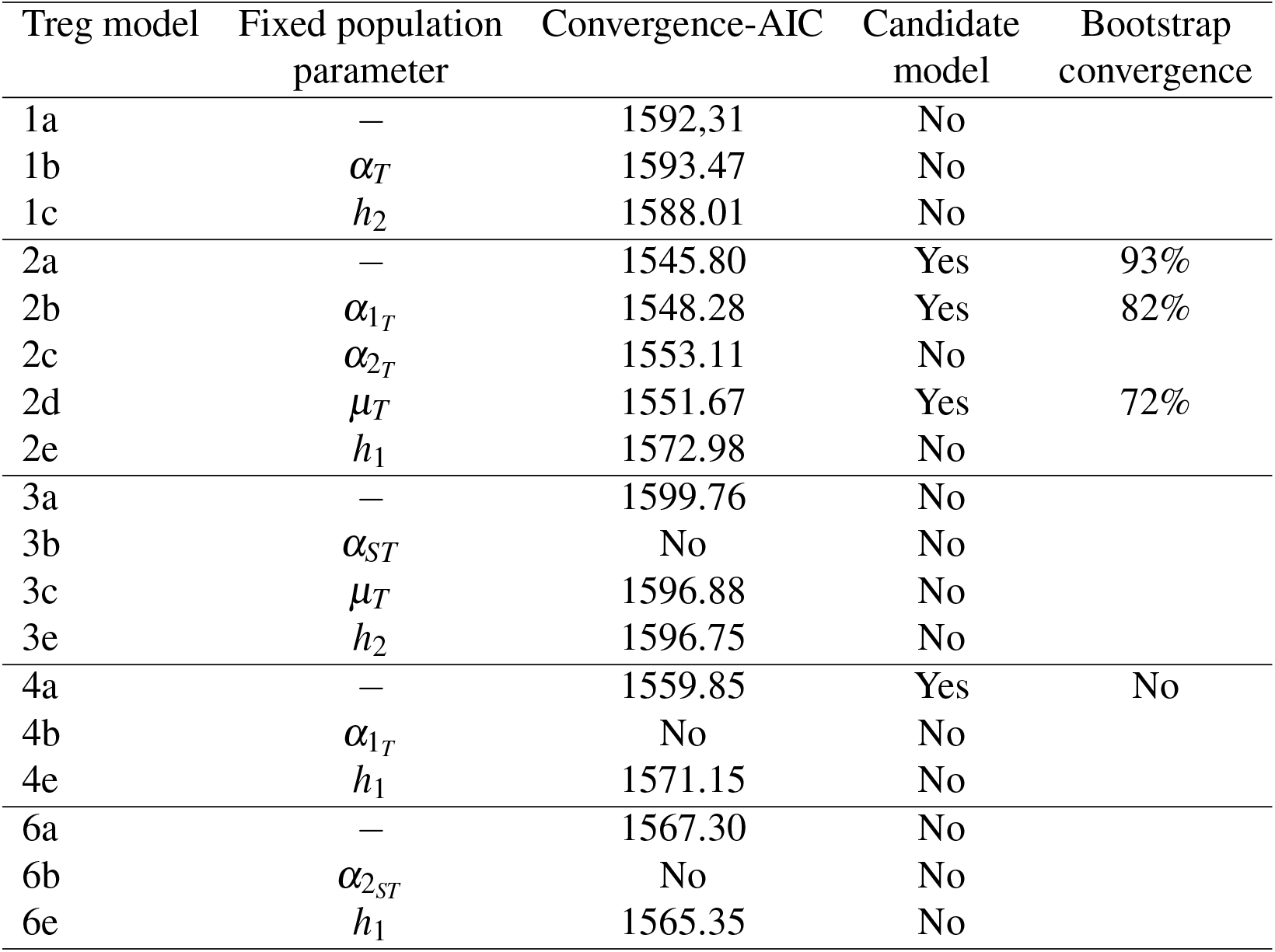
ODE Model formulations considered for memory conventional T-cell data and model selection procedure. Model 5 has been omitted from the table because non-convergence was obtained for random or fixed parameters.

| Treg model | Fixed population parameter | Convergence-AIC | Candidate model | Bootstrap convergence |
| --- | --- | --- | --- | --- |
| 1a | — | 1592.31 | No |  |
| 1b | $\alpha_T$ | 1593.47 | No | |
| 1c | $h_2$ | 1588.01 | No | |
| 2a | — | 1545.80 | Yes | 93% |
| 2b | $\alpha_{1T}$ | 1548.28 | Yes | 82% |
| 2c | $\alpha_{2T}$ | 1553.11 | No | |
| 2d | $\mu_T$ | 1551.67 | Yes | 72% |
| 2e | $h_1$ | 1572.98 | No | |
| 3a | — | 1599.76 | No |  |
| 3b | $\alpha_{ST}$ | No | No | |
| 3c | $\mu_T$ | 1596.88 | No | |
| 3e | $h_2$ | 1596.75 | No | |
| 4a | — | 1559.85 | Yes | No |
| 4b | $\alpha_{1T}$ | No | No | |
| 4e | $h_1$ | 1571.15 | No | |
| 6a | — | 1567.30 | No |  |
| 6b | $\alpha_{2ST}$ | No | No | |
| 6e | $h_1$ | 1565.35 | No | |

### 3.1 Conventional memory T-cells datasets (Tconv)

#### 3.1.1 Model selection

The Tconv dataset was modeled using the model selection process described in section 2.4.2. Initially, we considered model 1 for Tconv, which assumes 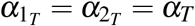. Model 1a supposed that all parameters had random effects, resulting in an AIC value of 1727.23.

We then considered model 2, where 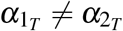. Model 2a assumed that all parameters had random effects, resulting in a converged AIC value of 1705.11. When we introduced fixed effects for *h*_2_, the model demonstrated a slightly lower AIC value of 1701.86 in model 2b.

Next, we assumed the distinction between short-lived and long-lived T cells (ST and LT). Model 3 assumed all parameters had random effects, while models 3b and 3c fixed *u*_*ST*_ and *h*_2_, respectively. Only model 3a, with an AIC value of 1756.3, showed convergence.

Model 4 considered different expansion rates of T cells after each vaccination. Model 4a considered random effects for all parameters, while models 4b and 4c fixed the expansion rates 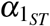 and 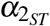. Model 4d assumed the decay of T cells (*μ*_*ST*_) occurred with a fixed population parameter. Models 4e and 4f fixed the period after each vaccination that T cells were activated for *h*_1_ and *h*_2_, respectively. SAEM convergence was only reached for model 4a and model 4c, with AIC values of 1651.33 and 1649.52, respectively.

Models 5 and 6 were obtained when LT activation with a constant proliferation rate was assumed. Model 5 assumed that the activation rates of ST were identical following each vaccination. This led to model 5a, with an AIC value of 1760.13, assuming all parameters were random. In model 5b, we assumed *h*_2_ to be fixed population parameters, showing an AIC value of 1758.05. We did not achieve SAEM convergence in models with fixed population parameters *u*_*ST*_, *h*_1_, and *α*_*LT*_.

The last model examined was model 6, in which different activation rates for ST were considered. We set all parameters as random parameters, leading to model 6a, with SAEM convergence reached and an AIC value of 1660.01. We set the decay rate of ST-cells (*u*_*ST*_), proliferation rate of LT (*a*_*LT*_), and activation periods (*h*_1_ and *h*_2_) as fixed parameters in models 6b, 6c, 6d, and 6e, respectively. We considered combinations of these fixed parameters but did not observe any improvement except for model 6e, with SAEM convergence achieved and an AIC value of 1656.23.

Model 4c was initially chosen as the leading candidate model due to its lowest AIC value of 1649.52. However, it was later rejected as a result of a 1000-sample bootstrap that failed to converge. Similarly, Models 6a and 6e also lacked proper bootstrap convergence. As a result, Model 4a

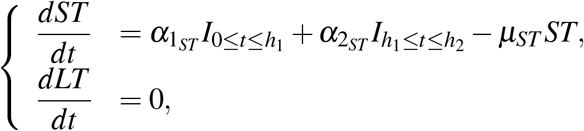

with an AIC value of 1651.33, was selected as a candidate model since it demonstrated bootstrap convergence with 95% of the bootstrap samples achieving SAEM convergence. A thorough search was conducted in both the converging and non-converging bootstrap datasets for frequently deviant profiles, but none were found. In summary, Model 4a indicates that the expansion rates of Tconv after the first and second vaccinations are distinct, and this may have implications for the long-term durability of vaccine-induced immunity. In addition, this model suggests the existence of two types of conventional memory T cells with one actively expanding and contracting after vaccination and the other remaining at a stable background equilibrium.

#### 3.1.2 Covariate influence

On the selected model 4a, it is now possible to investigate the influence of some covariates. To decide which parameter-covariate relationship to test next, we used the Conditional Sampling usage for Stepwise Approach based on Correlation testing (COSSAC)^22^ method that makes use of the data in the current model. By applying this method, the number of covariate models that are looked at is drastically reduced while the models that increase log-likelihood remain in the search. In particular, we investigated whether sex, age, antibody titres, TCR, early/late-converters, CMV, EBV and HSV seropositivity affect the model parameters.

The investigation revealed that the model examining early/late-converters and HSV seropositivity as a covariate, adding the effect on the parameter *μ*_*ST*_, generated a lower AIC of 1642.45, improving the AIC value of the original model selected by 8.68 points. The other covariates did not have a significant influence on model 4a. This result indicates that early/late-converters and HSV seropositivity may play an important role in the model. These results indicate that HSV1 carriership may be associated with lower expansion rates compared to non-carriers. Furthermore, early converters exhibit lower decay rates *μ*_*ST*_. The estimated parameter values and corresponding 95% confidence intervals for the final model 4a are shown in Table 4. Additionally, Figures 5 and 6 show the comparison between the observations of antibodies and the predictions from model 4a, as well as the Visual Predictive Check (VPC), which demonstrates that the observed percentiles match the expected percentiles and remain within the prediction intervals.

**Table 4.** Parameter estimates and corresponding 95% confidence intervals (CI) of final model 2a.

| Parameter | Estimate | 95% CI | P-value |
| --- | --- | --- | --- |
| $T_{reg}(0)$ | 13.13 | (0.812;27.418 ) | |
| $\alpha_{1T}$ | 7062.06 | ( 612.503;296344.9) | |
| $\alpha_{2T}$ | 4.3 | (0.769;11.013 ) | |
| $\beta_{\alpha_{2ST}}(EBV)$ | -1.4 | (-3.801;-0.163) | <2.2e-16 |
| $u_{ST}$ | 0.038 | (0.009, 0.083) | |
| $\beta_{u_{ST}}(Late - converter)$ | 1.76 | (1.105, 2.985) | 1.87e-14 |

**Figure 5.**
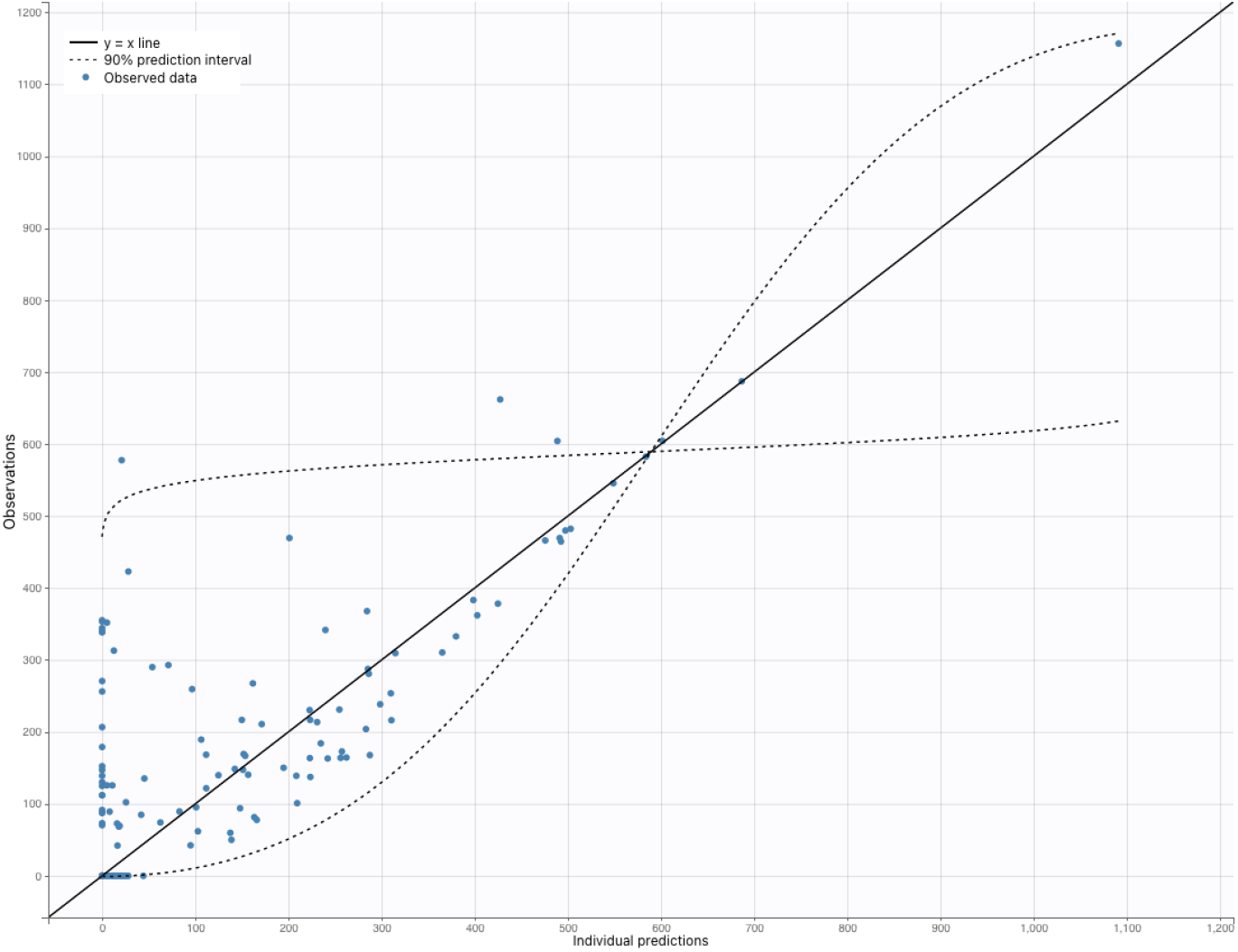
Comparison of observed memory conventional T-cell data (blue circles) with the predictions of model 4a (line). The shaded region around the red line represents the 95% confidence interval of the model predictions.

**Figure 6.**
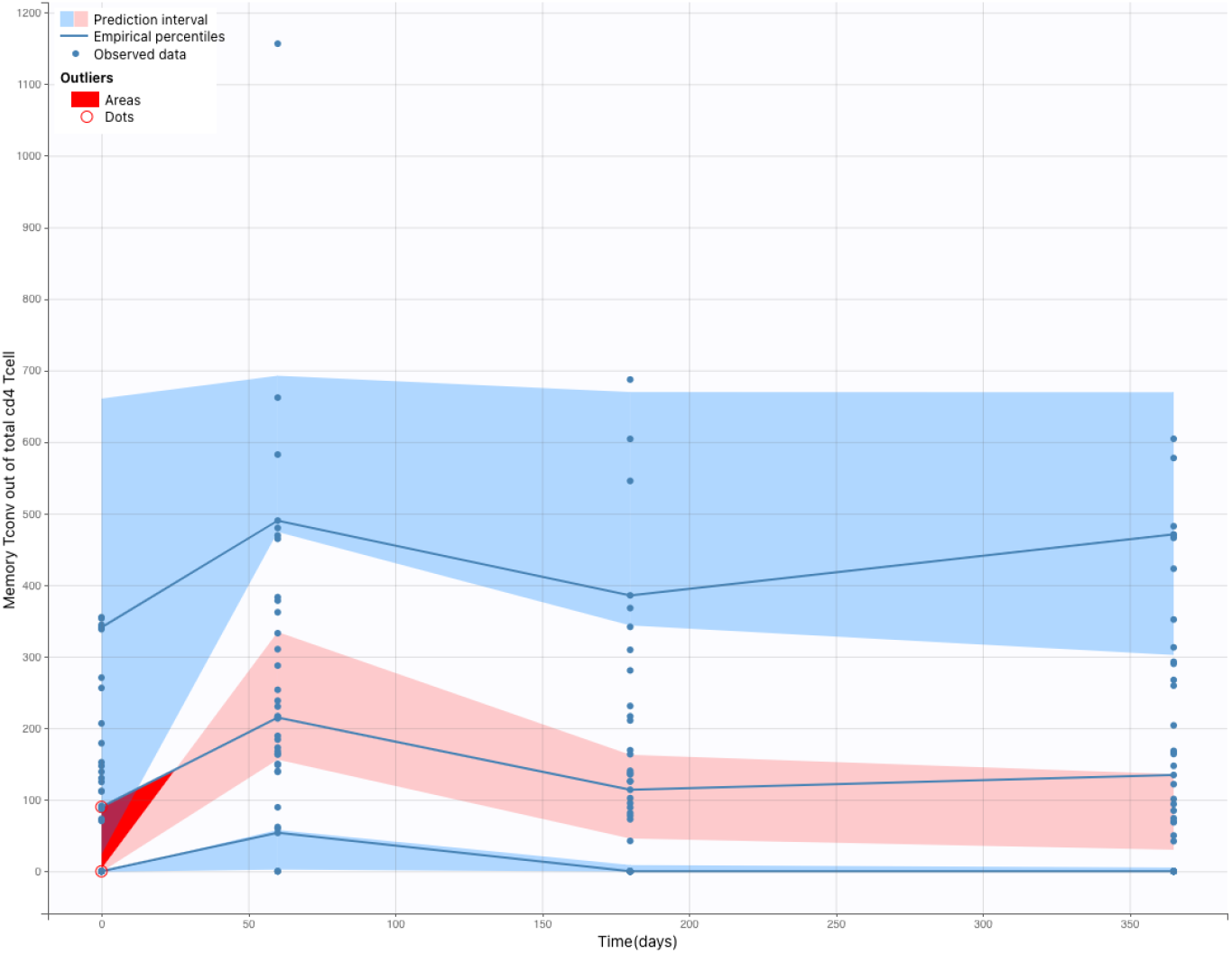
The Visual Predictive Check (VPC). The blue lines represent empirical percentiles that summarize the Tconv data. The blue and pink areas represent 95% prediction intervals and summarize predictions from the model 4a. The observed percentiles match the expected percentiles and remain within the prediction intervals.

### 3.2 Regulatory memory T-cells datasets (Treg)

#### 3.2.1 Model selection

Similarly as previously regarding the conventional memory T cells, we used model 1 to analyze the Treg dataset and observed the following distinctions: a scenario where all parameters have random effects led to model 1a, where an AIC value of 1592.31 was found. Setting *α*_*T*_ fixed in model 1b did not improve the model (AIC:1593.47). On the other hand, when we set fix *μ*_*T*_, *h*_1_ and *h*_2_ we get a lower AIC value for model 1c (AIC:1588.01), 1d (AIC:1590.1), 1e (AIC:1589.35), respectively.

Next, we considered model 2, which assumes a changed rate after the second vaccination due to a memory response. We obtained an AIC value of 1553.86 for model 2a, assuming all parameters had random effects. However, when we adapted the assumptions and considered fixed population parameters, the model did not improve in some cases and did not converge in others.

We further explored models that separated SB and LB. For model 3a, where all parameters were assumed to have random effects, we obtained an AIC value of 1599.76. However, no SAEM convergence was achieved for model 3b, where we fixed *α*_*T*_. In models 3c, 3d, and 3e, we fixed *mu*_*T*_, *h*_1_, and *h*_2_, respectively. Model 3c and 3e showed a lower AIC value, 1596.88 and 1596.75, respectively.

For model 4a, where all parameters were considered random, we obtained an AIC value of 1559.85. When we set 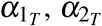, and *μ*_*ST*_ as fixed parameters in models 4b, 4c, and 4d, respectively, we did not achieve SAEM convergence. No significant result was obtained when setting other parameters fixed.

We also considered models 5 and 6, which include a proliferation rate for LT. For model 5, many assumptions were made about the parameters, but we did not obtain convergence. Finally, we examined model 6. When testing different assumptions about the parameters, only model 6a and model 6f showed convergence. In model 6a, all parameters had random effects, leading to an AIC value of 1567.3. In model 6f, *h*_2_ was set as a fixed parameter, giving an AIC value of 1564.35.

The model 2a had the lowest AIC value of 1545.8 among all the above-mentioned models

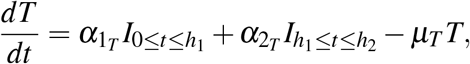

and was selected as the first candidate model. The candidate model showed bootstrap convergence; 93% of the bootstrap samples achieved SAEM convergence. As before, an examination of often divergent profiles in the converging and non-converging bootstrap datasets, but no such profile was detected.This suggests that the model is robust and that the results are reliable. Overall, these results indicate that model 2a is a strong candidate for explaining the underlying data and that it can be used to make predictions or draw conclusions about the phenomenon being studied. We note that model 2a is similar to model 4a with the difference of no co-existence of a stable background population.

#### 3.2.2 Covariate influence

We can identify the impact of the covariate on the candidate model 2a by using the COSSAC approach^22^. The model that examined early/late converters, adding the effect on the parameter *μ*_*ST*_, and EBV seropositivity, adding the effect on the initial value of *α*_2*T*_, was discovered to have the lowest AIC of 1536.89, decreasing the AIC value to the original model chosen by approximately 9 points. The estimated parameters and their 95% confidence intervals for the final Model 2a, which includes the effect of early/late converters and EBV seropositivity on the initial values of *μ*_*ST*_ and *α*_2*T*_, respectively, are shown in Table 4. Figure 8 shows the observations of antibodies compared to the predictions of model 2a, and the visual predictive check (VPC) comparing the model-based simulations with observed data. The VPC plot shows that the observed percentiles are close to the predicted percentiles and remain within the corresponding 95% prediction intervals, indicating that the model can accurately predict the immune response.

**Figure 7.**
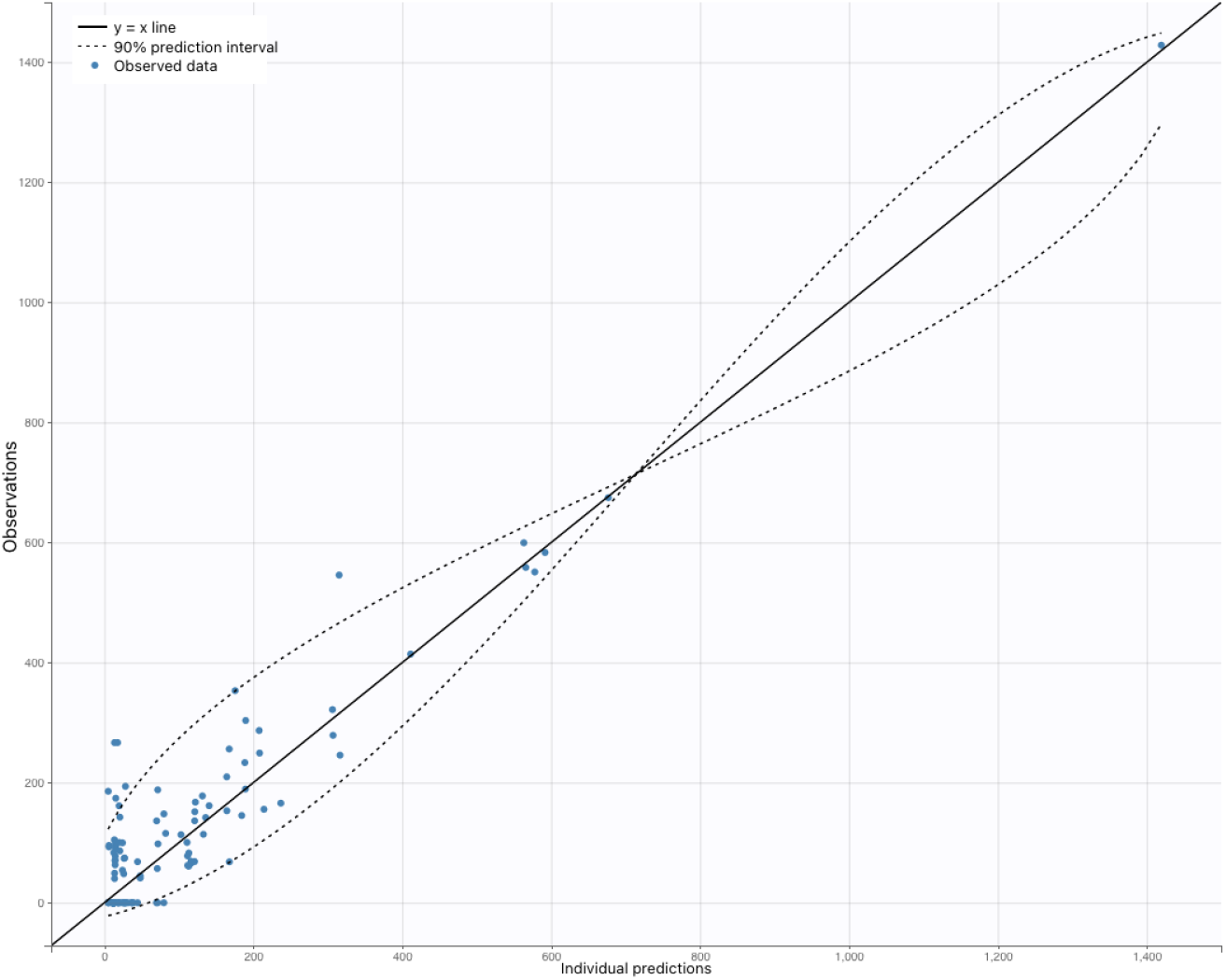
Comparison of observed memory regulatory T-cell data (blue circles) with the predictions of model 2a (line). The shaded region around the red line represents the 95% confidence interval of the model predictions.

**Figure 8.**
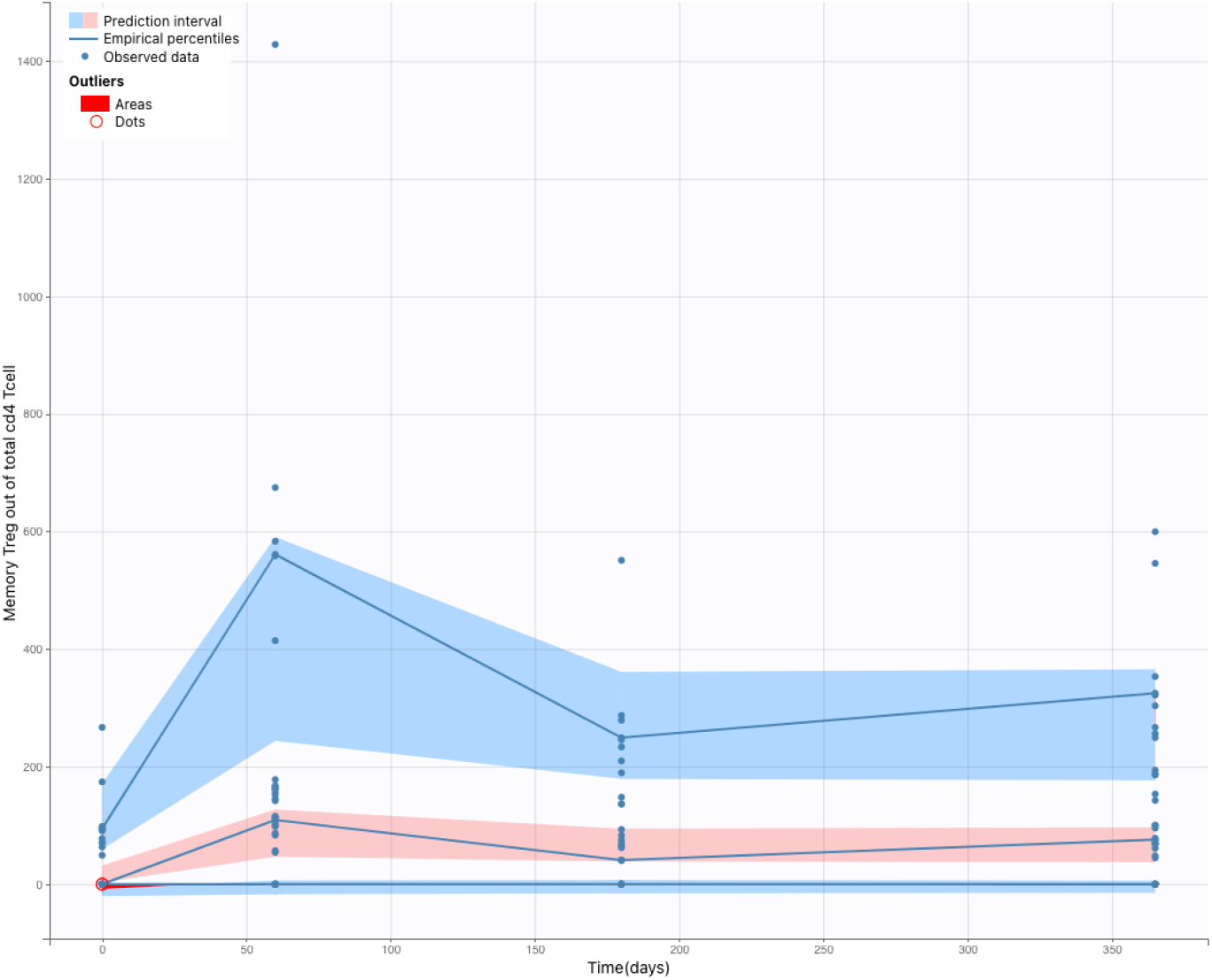
The Visual Predictive Check (VPC) comparing the results of the model-based simulations with observed data. The blue lines are empirical percentiles and summarize the observed data. The blue and pink areas are 95% prediction intervals and summarize predictions from the model 2a. The observed percentiles are close to the predicted percentiles and remain within the corresponding prediction intervals.

The results suggests that individuals who have been previously infected with EBV may have a different immune response to vaccination compared to those who have not been infected. Specifically, our study found that EBV seropositivity was associated with an increase in Treg cell expansion rates following vaccination, potentially enabling the virus to establish a stable infection by persisting within an individual.

Furthermore, the results also suggest that the dynamics of Treg cells may be influenced by the timing of their conversion. Specifically, individuals who convert to a positive response to the vaccine earlier have lower decay rates of Treg cells compared to late converters. We note that this finding is similar to what we found for Tconv.

## 4 Discussion and conclusion

Understanding the establishment of effective immunological memory is a complex task. In this study, we applied a mixed effects modeling approach based on ordinary differential equations to investigate the kinetics of hepatitis B virus surface antigen-specific CD4 T cells after vaccination and identified two models that contributed to the establishment of T cell memory. We modelled data available from a previously published vaccination trial with de novo Hepatitis B surface antigen^12^. In the previous paper the existence was described of early antibody producers (Early) and late antibody producers (Late).

i. Regarding conventional T cells, our best model (referred to as model 4) demonstrated that the dynamics of Tconv consist of two types of memory T-cells with actively expanding and contracting and the other remaining unchanged at a stable level. Moreover the dynamic Tconv type was influenced by the Early/Late converter status and HSV-1 seropositivity. It was observed that early converter vaccinees had lower decay rates for short lived Tconv cells, compared to late converter vaccinees and non-converters. This is supported by the finding in the Elias et al^12^ study, indicating that early-converters have a higher relative frequency of vaccine-specific TCR*β* sequences present in their conventional memory CD4 T cell repertoire at day 60 compared to vaccinees from the two other groups in the cohort. Our modeling results thus allow us to add knowledge on why these higher T-cell frequencies could have been established on day 60. In contrast to a potentially more rapid T-cell expansion in early converters, our modeling actually indicated that the higher frequency in early converters could be a consequence of a lower T cell decay rate in early converters compared to the two other converter groups. It was also revealed that the expansion rate after the second vaccination of the short lived Tconv cells was influenced by HSV-1 seropositivity, where the vaccinees without HSV-1 have a significantly higher expansion rate) compared to vaccinee who have HSV-1). Although other human herpesviruses have been noted to affect T cell responses upon vaccination (like CMV^23^), HSV-1 has not yet been reported by previous research. This could be suggesting that HSV-1 has the potential to modulate T-cell viability. Interestingly, HSV-infected cells have been reported to resist T-cell-induced apoptosis^24^, which may be a mechanism behind the observed effect.
ii. Regarding regulatory T cells, our best model (referred to as model 2) indicated no existence of a stable unchanged second Treg type next to the expanding and contracting Treg type. Our modeling indicated that the Treg expansion rate after the second vaccination for individuals with positive EBV seropositivity was higher than for those without it, thereby suggesting that EBV might induce Treg activity. Tregs are a type of immune cell that play a role in regulating the immune response and preventing autoimmune reactions. However, if their activity is too high, they may suppress the immune response to viral infections. This, in turn, could contribute to a higher level of virus in the body over time and facilitate the establishment and maintenance of viral persistence for EBV. This is consistent with previous studies in the literature^25–27^. Additionally, our analysis showed that the time it took for individuals to convert to a positive response to the vaccine (i.e., early converters vs. late converters) was associated with differences in Treg cell decay rates. Specifically, individuals who convert to a positive response to the vaccine earlier have lower decay rates of Treg cells compared to late converters. Hypothetically, like in Tconv, better TCR-epitope recognition could have rendered a lower Treg decay rate, but this hypothesis would only be valid if the TCR-epitope recognition for Tconv would be correlated with Treg cellular regulation, which still remains to be proven. However, We found a slightly significant effect of TCR on the decay rate of Treg cells in our analysis, but due to lack of data for some individuals and instability in the bootstrap, we were unable to include it in our final model. It is possible that differences in TCR could still influence Treg cell activity in a subtle or complex manner, and further research is warranted to better understand its role in Treg cell dynamics.

In summary, our mixed effects modeling approach, based on ordinary differential equations, identified key factors influencing effective immunological memory in Tconv and Treg. Our study provides novel quantitative evaluation of T-cell dynamics temporal scales, offering insights into crucial biological processes unattainable with traditional statistics. We incorporated phenomenological elements to capture trends and interindividual variability. By combining mechanistic and phenomenological components, our models comprehensively elucidate kinetics and factors driving T cell memory establishment.

This integration of mechanistic and phenomenological models enhances our understanding of immunological memory mechanisms after vaccination, bridging the gap between detailed insights and predictive capabilities. Our study’s value and applicability are heightened by this approach, supporting more comprehensive and nuanced research in quantitative immunology. Moreover, this knowledge directly contributes to infectious disease immunology and establishes a solid foundation for further quantitative immunology research.

Furthermore, in the field of vaccination,our findings offer insights that contribute to a better understanding of the long-term dynamics of vaccine-induced immunity. By studying immune cell dynamics and identifying key factors that influence sustained immune protection, we can gain knowledge that helps in predicting how the immune response will develop over time. This understanding is valuable for clinical trial researchers as it enables them to make more informed assessments of vaccine effectiveness and design optimal vaccination strategies. It also aids in identifying individuals who may benefit from additional interventions to maintain their immune response. Overall, our findings contribute to advancing our knowledge of vaccine-induced immunity and support evidence-based decision-making in the field of vaccination.

However, it is important to acknowledge the limitations of our research. Firstly, our study focused primarily on Treg and Tconv cells, and did not extensively explore other immune cell populations such as B cells. Additionally, our modeling approach, while informative, simplifies the complexity of the immune system and may not capture all aspects of immune dynamics. Furthermore, our analysis relied on available data, which had its own limitations in terms of sample size and data quality. Future studies should aim to address these limitations by incorporating a broader range of immune cell types and more comprehensive datasets.

In conclusion, our study has provided valuable insights into the dynamics of regulatory T cells (Tregs) and conventional T cells (Tconv), highlighting their distinct responses to various factors. These findings underscore the importance of considering the diversity within the immune system. Moving forward, it is crucial to validate and refine our models using comprehensive data, including longitudinal measurements of different variables. By advancing our understanding of immune dynamics, we can enhance our ability to develop precise and effective strategies to address immunological challenges.

## Acknowledgements

We would like to express our gratitude to Geert Mortier and Viggo Van Tendeloo for their valuable contributions and assistance in this research.

## Funding

BO acknowledges funding received from the European Research Council (ERC) under the European Union’s Horizon 2020 research and innovation programme (grant agreement 851752-CELLULO-EPI) and Research Foundation Flanders (FWO) (grant agreement 1861219N). PB and PVD received UAntwerp funding (Methusalem and concerted research action).

## Notes

### Competing Interest Statement

BO declares that the University Hospital Antwerpen and the University of Antwerp have received investigator-initiated grants from Abbvie, Pfizer and Roche. BO is a shareholder and member of the board of ImmuneWatch BV. NH declares that the Universities of Antwerp and Hasselt have received funding for advisory boards and research projects of MSD, GSK, JnJ, Pfizer and the European Commission's IMI programme outside the proposed work. NH has not received any personal remuneration. PB declares the University of Antwerp received compensation for unrelated consultancy for Pfizer in 2019, research grants from MSD and Pfizer, and the European Commission's IMI programme. PB has not received any personal remuneration.
The remaining authors declare that the research was conducted in the absence of any commercial or financial relationships that could be construed as a potential conflict of interest.
KL is a shareholder and member of the board of ImmuneWatch BV.
PM is a shareholder, employee and member of the board of ImmuneWatch BV.

### Summary of Updates

In the revised version of the manuscript, a modification has been made pertaining to the authorship. Specifically, a request has been received from two individuals involved in the initial publication to have their names excluded from the author list.

